# Gene Expression Analysis of Glioma Neural Stem Cells Shows Disturbed Amino Acids Metabolism and Axonal Growth Cone Dynamics in Glioblastoma Multiforme

**DOI:** 10.1101/2021.09.27.461538

**Authors:** Rutvi Vaja

## Abstract

Glioblastoma multiforme(GBM) is a group of fatal and aggressive tumors of the central nervous system. Despite advancements in the treatment of GBM, patients diagnosed with these tumors typically have a poor prognosis and poor quality of life as the disease develops. The single-cell RNA high-throughput sequencing processed data for Glioma cancer stem cells were taken from GEO and analyzed to find out the underlying expression differences at the gene level between glioma neural stem cells(GSCs) and Normal neural stem cells(NSCs). In the current study, we have performed an RNA-sequencing analysis between GSCs and NSCs to better understand the origin of GBM. We have performed bioinformatics analysis on the transcriptional profile of 134 samples which consisted of 75 GSCs and 59 NSCs obtained from the NCBI bio project(PRJNA546254). First, an exploratory analysis was performed which showed significant variation patterns between GSCs and NSCs. Subsequently, Deseq2 differential gene expression analysis identified 1436 differentially expressed genes between GSCs and NSCs[(padj. value <0.05, log2 fold change (>=+/-1.5)]. This study reveals genes like *MAOA, MAOB, GATM, GLDC, AMT, and SHMT1* as the key features contributing to the disturbed processes of Glycine, threonine, and serine amino acid metabolism, axonal cone growth curve, and cell migration in Glioma. Conclusively, our study also depicts gene expression changes in amyloid beta-binding protein in between GSCs and NSCs which plays an important role in tumor microenvironment formation. Besides, the results presented here reveal new insight into the progression of GBM and the identification of novel genes involved in gliomagenesis.

## Introduction

Glioblastoma Multiforme(GBM) is the most common malignant brain tumor(1). GBM accounts for the most aggressive brain tumors arising from glial cells(2). It has a median survival rate of 12.6 months(3).GBM has widespread and infiltrative development into adjacent brain tissue, increased cell growth, the ability to overcome cell death and antitumor immune responses, and therapeutic resistance(4). GBM recurrence is linked to (i) radio- and chemotherapy resistance; (ii) diffuse features due to tumor cells’ invasiveness properties over the surrounding brain parenchyma; and (iii) tumor intra- and inter-heterogeneity(5). GBM has no risk factors other than rare instances of genetic predisposition and irradiation exposure(7).GBM most commonly occurs in the cerebral hemispheres, with 95 percent of tumors occurring in the supratentorial region, compared to only a few percent in the cerebellum, brainstem, and spinal cord(9).

WHO (World Health Organization) classification is the current international standard for glioma terminology and diagnosis. It divides gliomas into four grades(Grades 1 to 4) based on the degree of malignancy assessed by histological criteria(6). The standard treatments for GBM remain unchanged for many years-which involves surgical resection, chemotherapy, and radiotherapy, despite no significant goal being achieved in treating this horrendous disease(7). The prognosis of GBM is very poor, which makes it a crucial public health issue(8).

Many efforts have been made by researchers and oncologists to understand the origin of GBM, but the reason still remains unclear. Due to this, the etiology of GBM remains unknown. Out of the many hypotheses, two of them suggest the origin of GBM. One of them suggests that GBM arises from Cancer stem cells(CSCs) that possess the ability to self-renew, differentiate and encourage tumor formation. The second theory suggests that GBM arises from normal neural stem cells once they acquire several different types of mutations in common neural marker cell genes(10). This points to the importance of understanding the mechanism behind how neural stem cells transform to Glioma neural stem cells via acquired mutations. This can be achieved by understanding the gene expression differences between GSCs and NSCs.

The main goal of our study was to identify the underlying gene expression level changes between Glioma neural stem cells and Normal neural stem cells. Our main concern was to identify the affected biological pathways in Glioblastoma Multiforme which can be used as a potential tool for the cure of GBM. Identification of affected biological pathways would help further in the identification of molecular mechanisms which encourage CSCs to proliferate tumor formation. We wanted to find out potential key biomarkers for targeting CSCs which suggest the origin of GBM.

Hence, in the present study, we investigate the transcriptomic profiles of 75 Glioma Neural stem cells samples and 59 Normal neural stem cells samples to understand the gene expression differences between them. After the identification of significant differentially expressed genes, we scrutinized the most affected biological pathways which revealed important results into the neuronal processes like synaptogenesis and progenitor dynamics as the transformations happen from CSCs to GBM or NSCs to GBM. Besides, the results presented here reveal new insight into the progression of GBM and the identification of novel genes involved in gliomagenesis.

## METHODS

### Softwares used in this study

#### Server T-Bio Info

Website: https://server.t-bio.info/

The Principal component analysis, H-Clust. Heat Map plots and Differential gene expression analysis was done using this Cloud based server.

#### Metaboanalyst

Website: https://www.metaboanalyst.ca/

Metaboanalyst was used to generate PCA plots using the associated data set of this study

#### Enrichr

Website: https://maayanlab.cloud/Enrichr/

The Gene ontology analysis was performed using the Enrichr Knowledge database.

#### Data Sets

In this study, the process of single-cell RNA high-throughput sequencing processed data for Glioma cancer stem cells was taken from NCBI GEO[GSE132172]. The dataset of this project was generated by Zhao Y et. al(2019) and was published as a bio project on NCBI with accession number(PRJNA546254). This dataset consisted of RNA-Seq data retrieved from CB660 normal neural stem cell lines and GliNS2 glioblastoma stem cell lines. A total of 134 samples were present on the associated SRA Run Selector, 59 samples of NSCs, and 75 samples of GSCs were selected and downloaded as an SRA Run.

**Table 1:** Dataset used in this study.

| **Disease State** | **Number of samples** |
| --- | --- |
| Glioma Neural stem cells | 75 |
| Normal neural stem cells | 59 |

#### Data Pre-processing

The high throughput sequencing processed data was quantile normalized signal data. The processed data had gene symbols.

### Down-stream Analysis

#### Principal Component Analysis

To understand the patterns of gene expression in the data, comparative data analysis was performed using the Principal component analysis(PCA) module integrated on the T-bio info server (https://server.t-bio.info/). PCA is a dimensional reduction technique that is applied to larger data sets, in order to visualize the variation between samples in a particular data set(11). PCA was performed between these 2 conditions namely : (a) Glioma neural stem cells(GSCs) and (b) Normal neural stem cells(NSCs).Eventually, the PCA plots were generated using Metaboanalyst (https://www.metaboanalyst.ca/)(12).

#### Differential Gene Expression Analysis

The differential gene expression analysis was performed using the Deseq2(13) pipeline on the Tbio-info server to derive significantly differentially expressed genes between GSCs and NSCs samples. DESeq2 is a method for differential analysis of count data that uses shrinkage estimation for dispersions and fold changes to improve the stability and interpretability of estimates. This enables a more quantitative analysis focused on the strength rather than the mere presence of differential expression. The significant genes were identified with the threshold of (p.adj value <0.05, Fold change (>= ±1.5)(14).

#### Assessment of Discriminatory potential of Significant genes

In order to visualize and obtain the potential significant genes, H-clustering(15,16) and a heat map(17) were formed using the 134 samples(75 GSCs and 59 NSCs) with the selected set of significant RNA genes. H-clustering is an important technique of machine learning which is used to group similar data points such that the points in each same group are more similar to each other than the points in the other groups. The groups formed are known as “Clusters.” H-clustering (Distance: Euclidean, Linkage: average) was performed to understand the discriminatory potential of significant genes in distinguishing the gene expression patterns between GSCs and NSCs samples. Eventually, Heatmaps were drawn using the Tbio-info server in order to elucidate specific up-regulated and down-regulated genes out of the potential significant genes obtained using Deseq2.

#### Gene enrichment Analysis

In order to elucidate the biological significance of the differentially expressed genes, a gene set enrichment analysis(18) was performed. Gene set enrichment analysis(GSEA) is a powerful analytical method to interpret gene expression data(19). GSEA was performed using Enrichr Platform(20), which is an easy-to-use intuitive enrichment analysis web-based tool providing various types of visualization summaries of collective functions of gene lists. Further, Gene set enrichment analysis helped to understand the biological role of the significant genes in the process of gliomagenesis.

## RESULTS

In this study, we analyzed the transcriptomic profile and data of the Glioma patient’s neural stem cells with that of the normal patient’s neural stem cells, in order to identify gene expression differences between them. The complete workflow of the study is represented in Figure 3 below.

**Figure 1:**
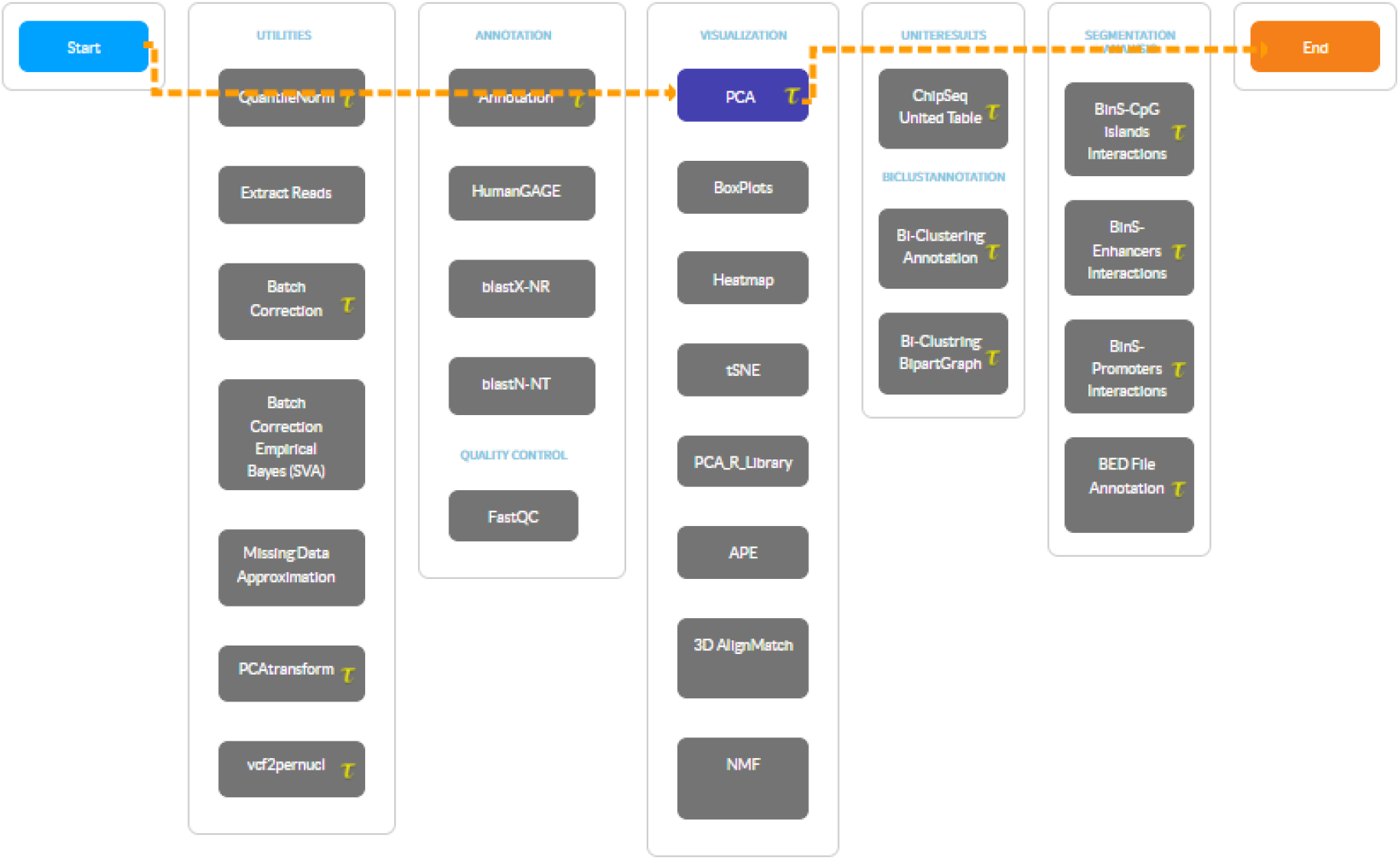
Screenshot of PCA Pipeline on T-bioinfo Server.

**Figure 2:**
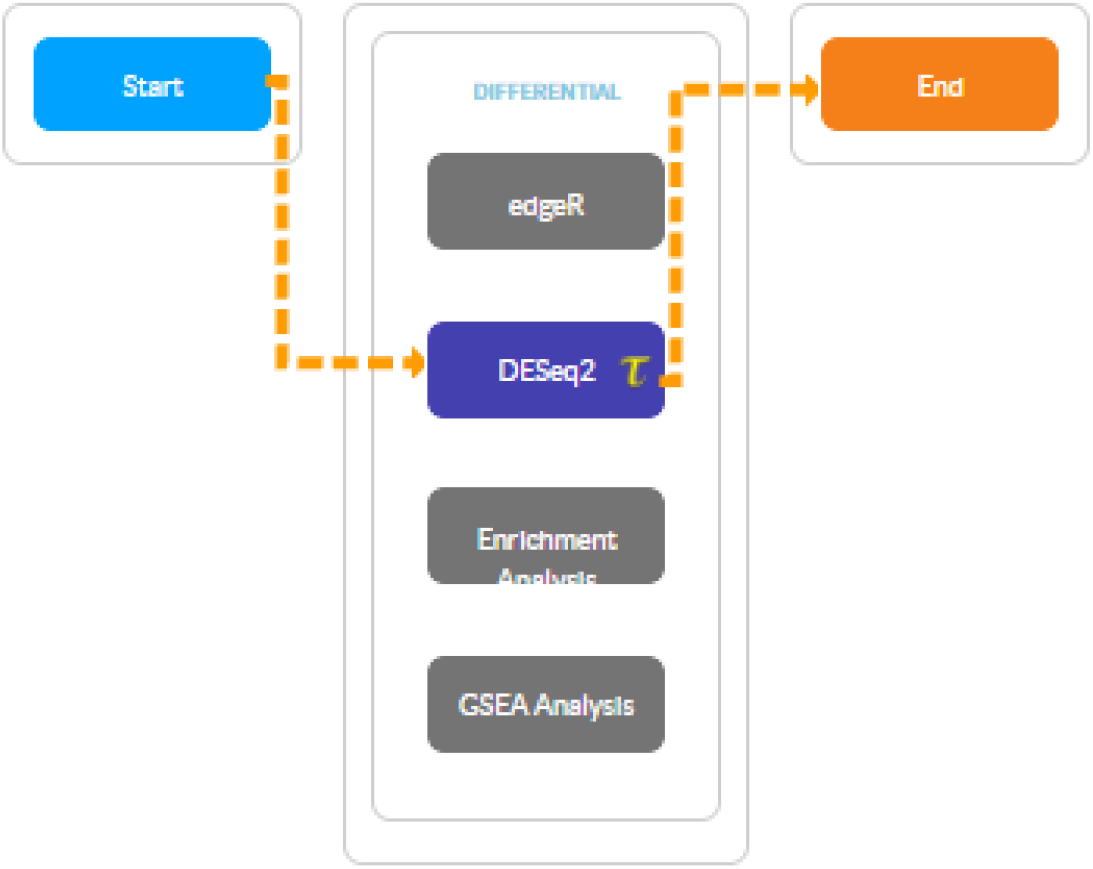
Deseq2 Pipeline for Differential gene expression Analysis on Tbio-info serve.

**Figure 3:**
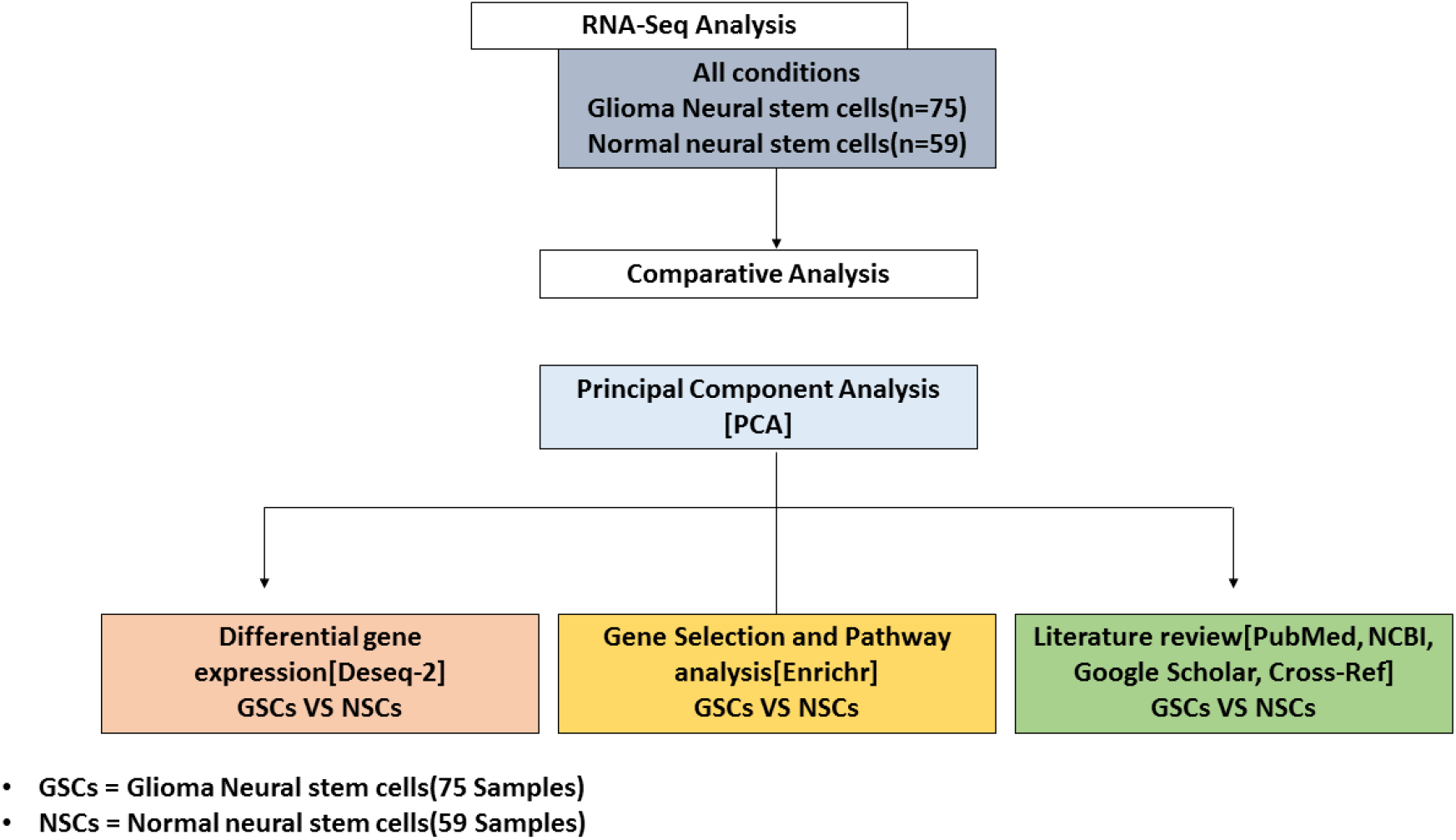
Work Flow of the study representing the key steps.

### Comparative Data Analysis

Data were analyzed using principal component analysis(PCA)(21) in order to elucidate the variation between these Glioma neural stem cells and normal neural stem cells. Figure 4 shows the variation between GSCs and NSCs. The PC1 is 14.30% and PC2 is 4.34%.

**Figure 4:**
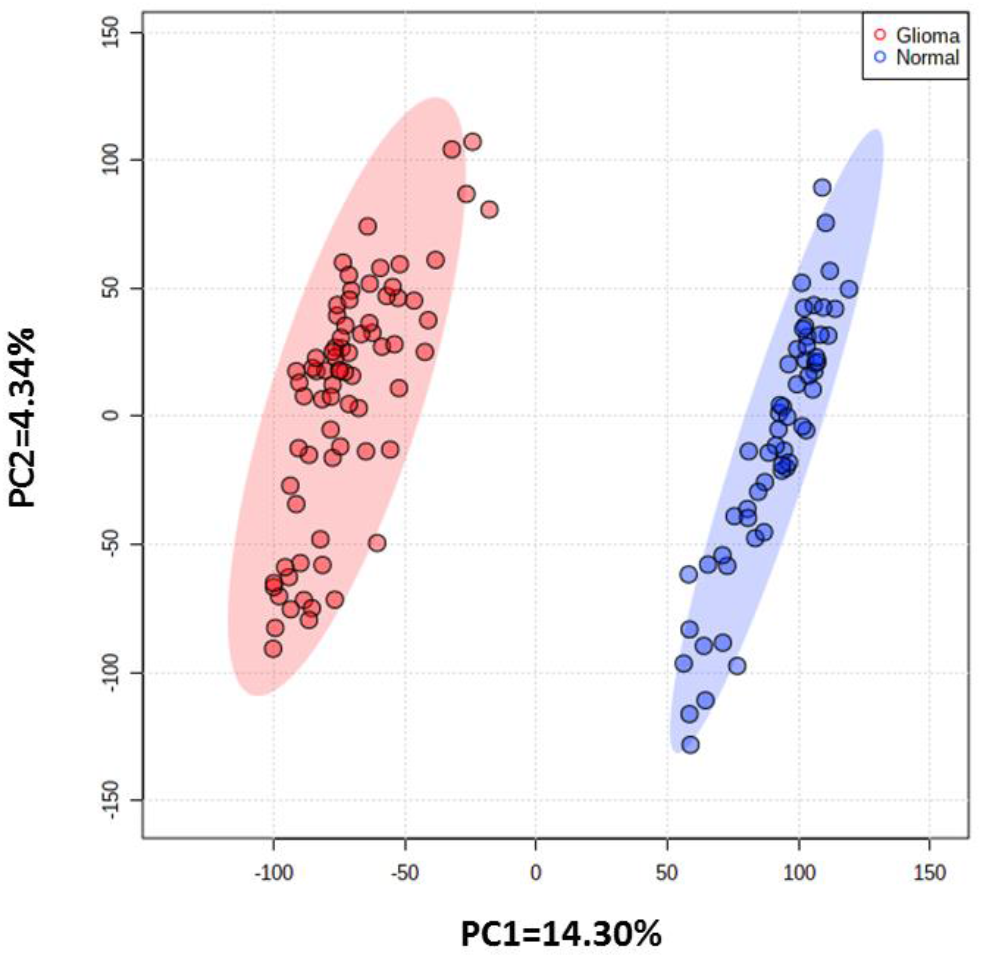
PCA between GSCs and NSCs samples.

### Downstream Analysis

It is evident from the PCA results Figure 4A that the variation between Glioma neural stem cells samples and Normal neural stem cells samples is maximum. Hence, we further carried out a comparative analysis between Glioma neural stem cells samples and Normal neural stem cell samples. Hence, in downstream analysis, we performed differential gene expression analysis between Glioma neural stem cells samples and Normal neural stem cell samples. Also, when the Glioma neural stem cells samples and Normal neural stem cell samples were examined in a specific scatter plot(22), it was obvious that there was an improvement in principal components, and the two groups were separated from each other which showed clear distinction. After the observed variation between these two groups-GSCs and NSCs, the reason for this difference and its impacts could be investigated at the transcriptomics level.

### Differential gene expression analysis

The differential gene expression analysis(23) between the Glioma neural stem cells(GSCs) and Normal neural stem cells(NSCs) samples scrutinized 1436 significantly differentially expressed genes between GSCs and NSCs[(padj. value <0.05, log2 fold change (>=+/-1.5)]. Among them, 619 genes were found to be significantly upregulated (p.adj value <0.05, Fold change >=+1.5) genes and 817 genes were found to be significantly downregulated (p.adj value <0.05, Fold change <= −1.5) in GSCs in comparison to NSCs.A volcano-plot for the differentially expressed genes can be seen in Supplementary Figure 1.

### Clustering and Heat Map revealed variations among Glioma neural stem cells and Normal neural stem cells samples

Hierarchical clustering(24) was performed in order to see clear variations between GSCs samples and NSCs samples. The main aim of hierarchical clustering was to understand if the significantly differentiated genes are capable of forming clusters based on their gene expression profiles. The clustering results show clear distinct variables between GSCs and NSCs samples as shown in Figure 5. The heatmap(25) representing the expression patterns between GSCs samples and NSCs samples is shown in Figure 6.

**Figure 5:**
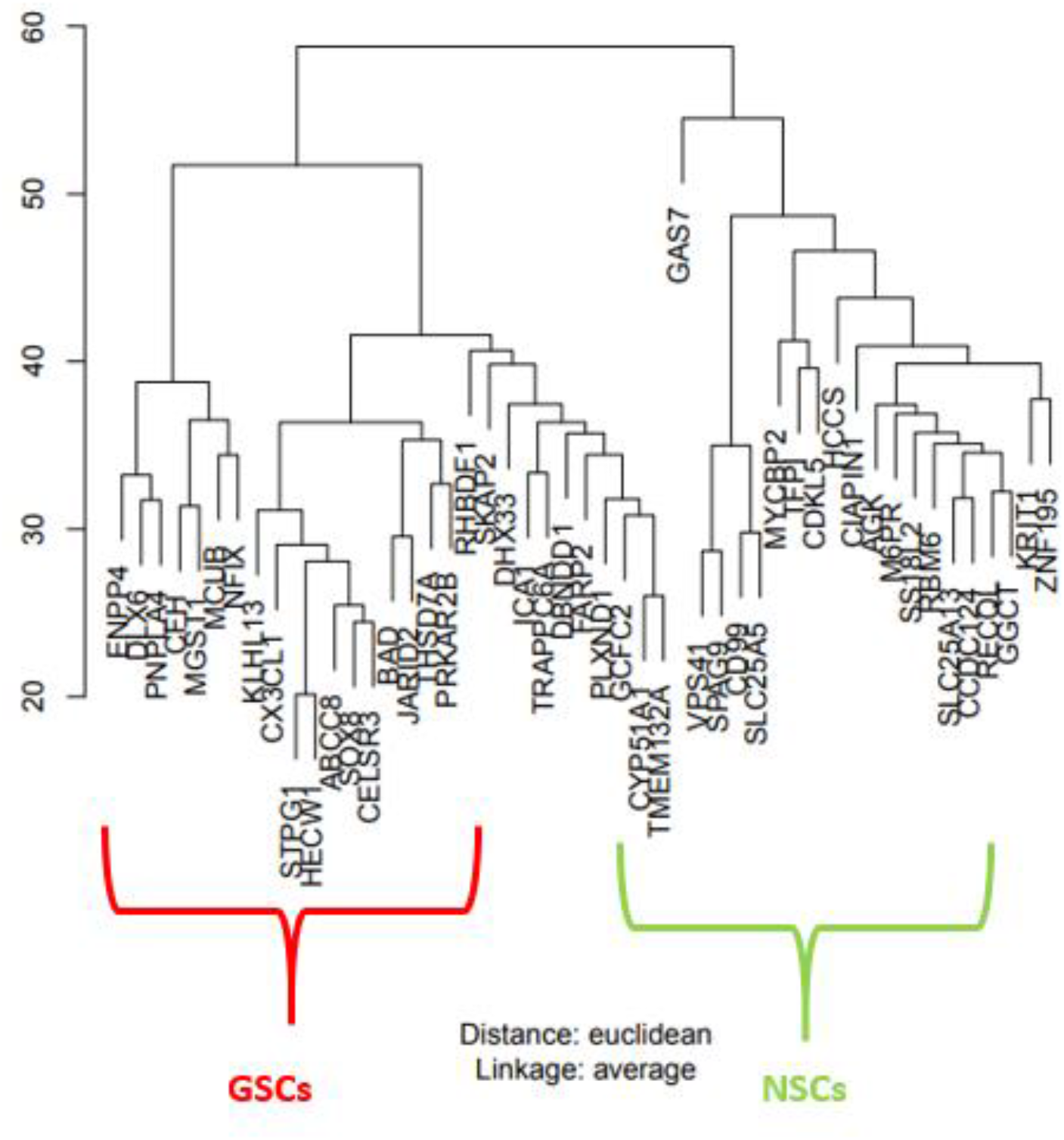
Hierarchical Clustering results as dendrograms. Red box clusters indicate the GSCs samples, and the green one represents the clusters of the NSCs samples.

**Figure 6:**
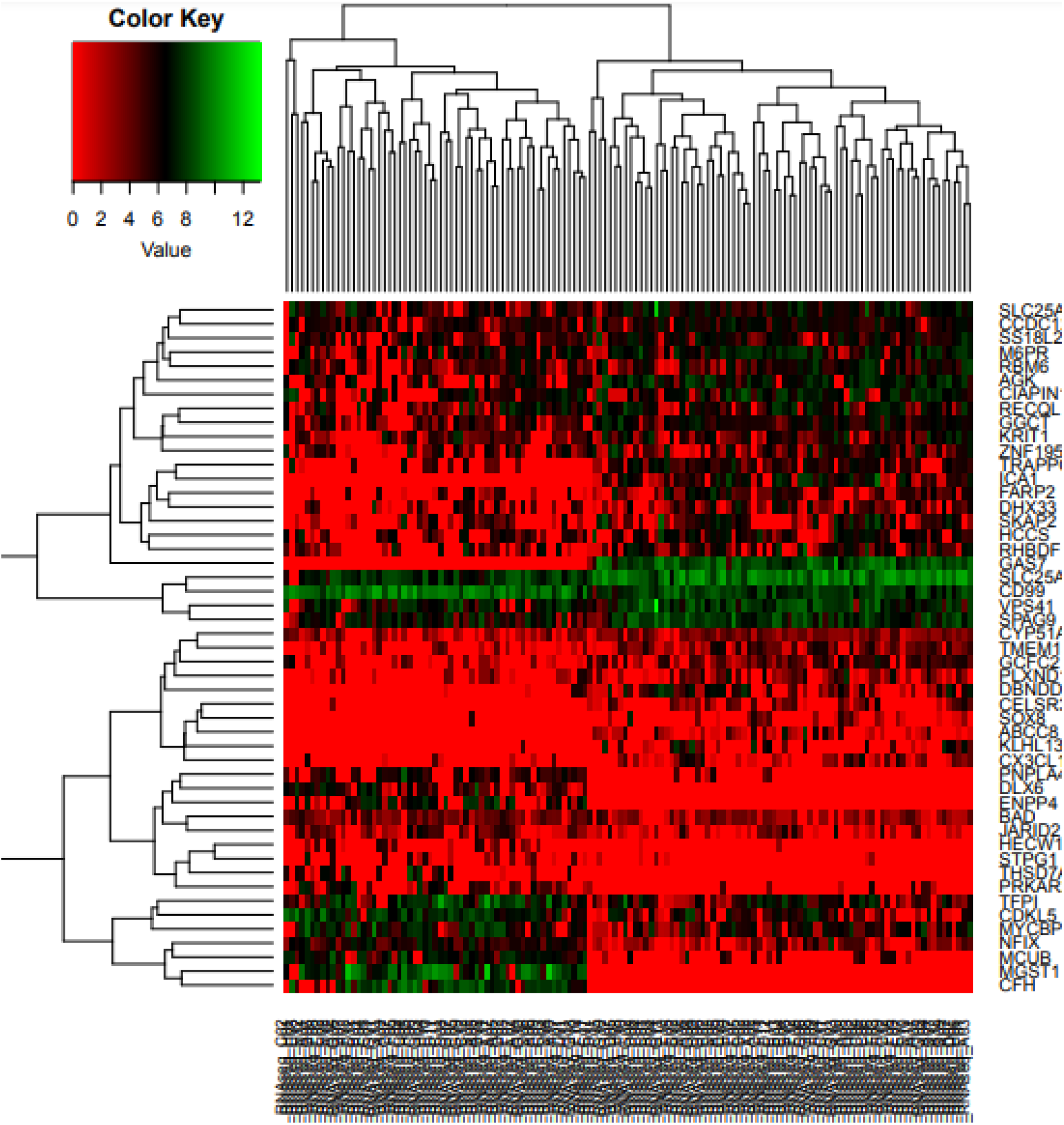
Heatmap representing the Gene expression patterns of significantly differentially expressed genes.

### Pathways involved in the pathogenesis of Glioma tumors by Gliomagenesis

To understand the involvement of the differentially expressed downregulated and up-regulated significant genes, pathway analysis(26) was performed using the Enrichr(27,28) knowledge database. Although the up-regulated genes showed involvement in varied types of pathways, some of the most important pathways were Axon guidance, Extracellular Matrix organization, Collagen binding pathways, and receptor-ligand binding pathways(Figure 7A).

**Fig 7A:**
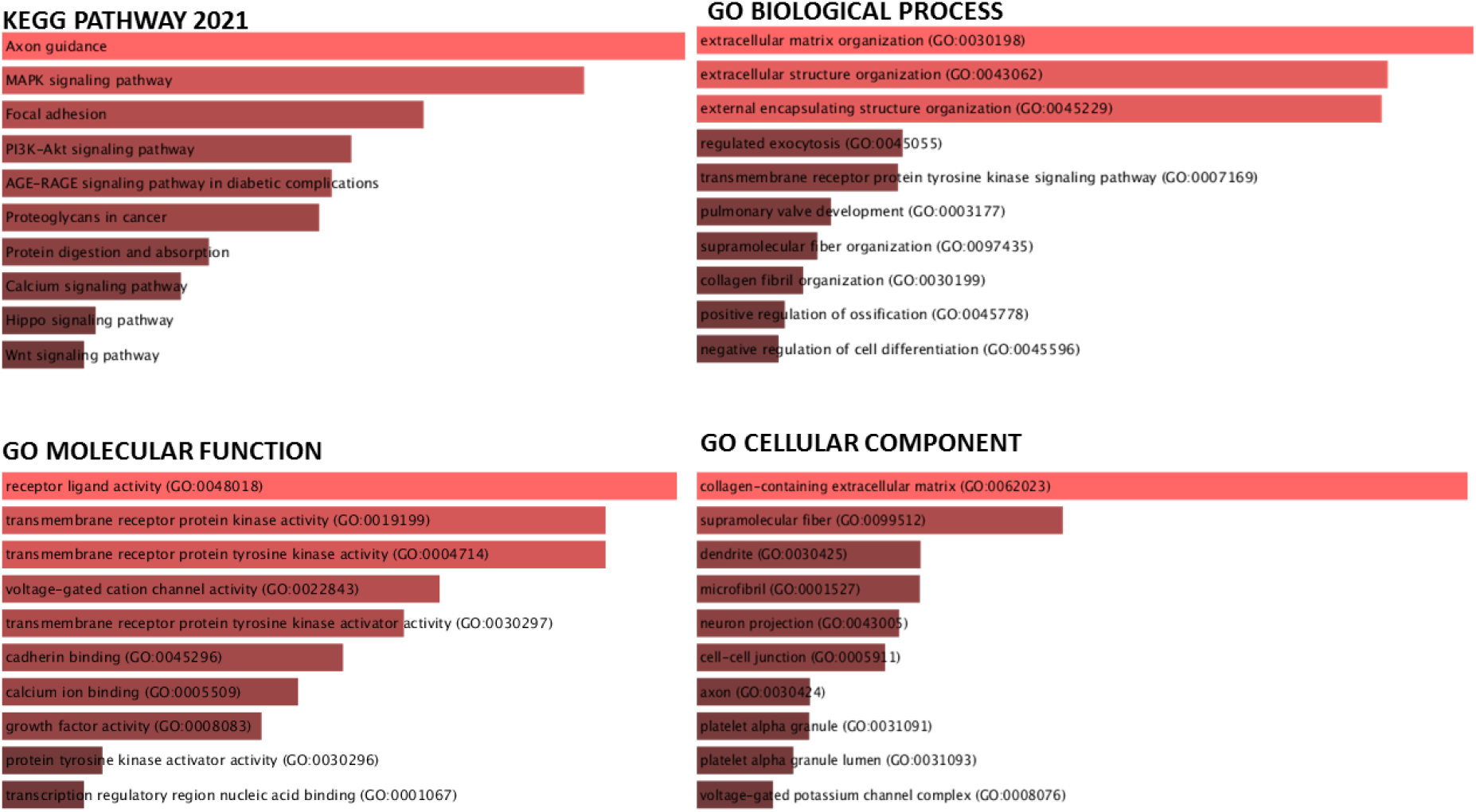
Gene ontology and KEGG pathway analysis of Up-regulated genes.

The same method was applied to find out the involvement of down-regulated genes with different biological pathways as shown in Figure 7B. The gene ontology analysis for down-regulated genes involved the Axon growth curve pathway, molecular nervous system development pathway, amyloid beta-binding pathways, and some of the amino acids metabolism pathway.

**Fig 7B:**
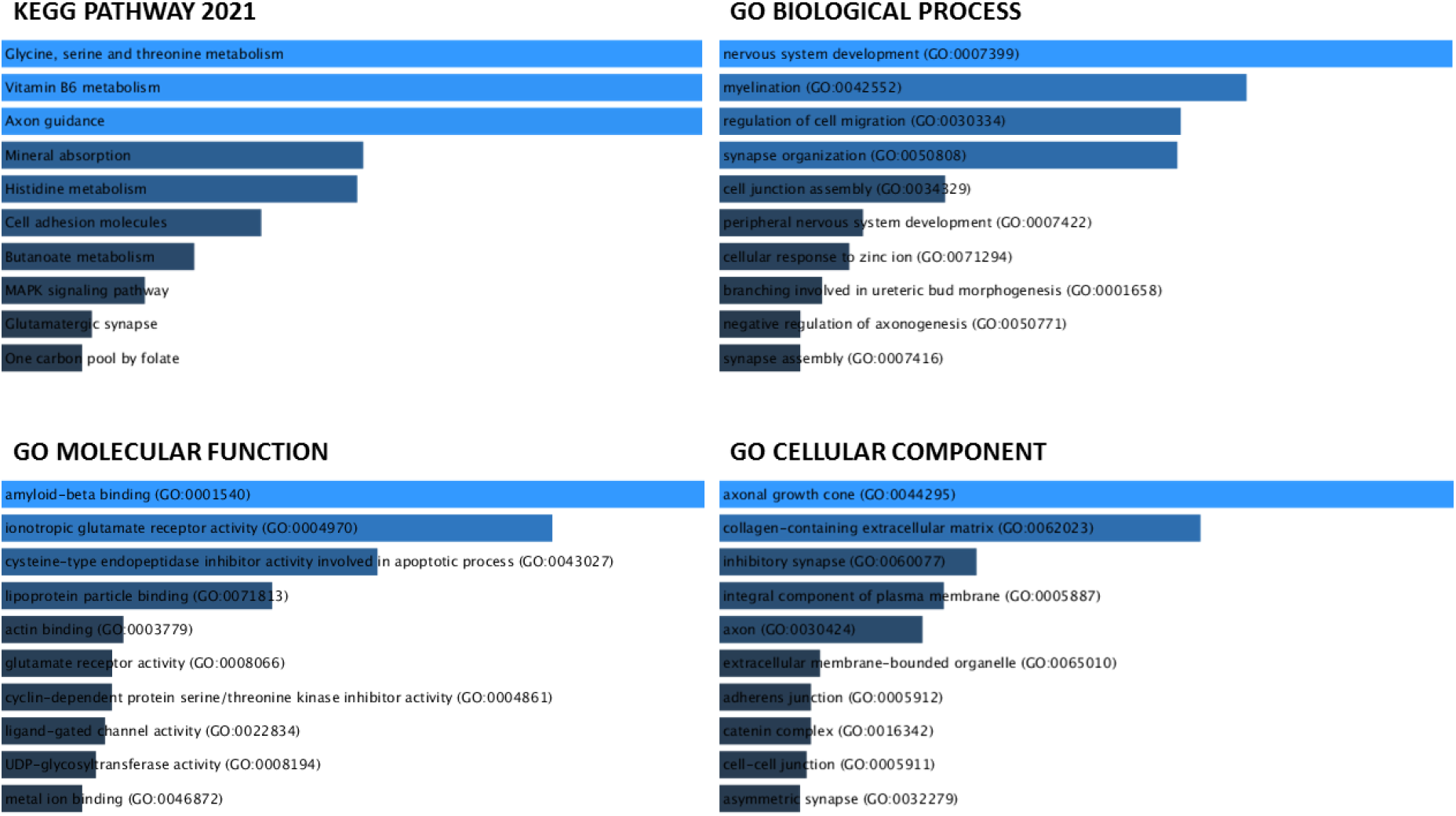
Gene ontology and KEGG pathway analysis of Down-regulated genes.

On the basis of Principal Component analysis, there is a clear distinction of groups formed between GSCs and NSCs samples which denotes variability. However, disruption of amino acid metabolism namely threonine, serine, and Glycine is what we focus our discussion and conclusions on. Interestingly, the disturbing biological processes of the axonal growth curve reveal many deeper insights herein into the process of gliomagenesis, which is also discussed further.

The Gene ontology results show the involvement of genes like *MAOA, MAOB, GATM, GLDC, AMT, and SHMT1* as the key features contributing to the disturbed processes of Glycine, threonine, and serine amino acid metabolism. Apart from this genes like *COBL, FLRT3 and GPM6A* contribute to the significantly downregulated disturbing process of the axonal growth curve in humans.

**Table 2.** Differentially Expressed 6 Genes identified in our study which are involved in disturbing Amino acid(Glycine, serine, and threonine) metabolism(p.adj value <0.05, Fold change <= −1.5).

| Sr. No. | Gene Symbol | Gene Description | P-adjusted Value |
| --- | --- | --- | --- |
| 1. | MAOA | monoamine oxidase<br>A | 1.71E-09 |
| 2. | MAOB | monoamine oxidase<br>B | 0.003507469 |
| 3. | GATM | glycine<br>amidinotransferase | 3.55E-28 |
| 4. | GLDC | glycine<br>decarboxylase | 0.019216624 |
| 5. | AMT | aminomethyltransfer<br>ase | 0.00017872 |
| 6. | SHMT1 | serine<br>hydroxymethyltransf<br>erase 1 | 8.55E-05 |

## Discussion

The origin of Glioma and progression of Gliomagenesis still remains unknown(29). With no known particular etiological aspect addressed to date, this calls for exploring further the role of glioma tumors. In this study using the RNA-Seq analysis approach on the transcriptomic profiles of Glioma and Normal neural stem cells, we have tried to establish a close connection between Glioma tumor formation and the underlying gene expression changes. Comparative data analysis using PCA revealed that the samples of both categories form separate clusters, depicting the vast genetic differences between the GSCs and NSCs samples. This drives a common theory that there is something at the transcriptomics level that leads to the variation between Glioma neural stem cells and Normal neural stem cells. Further studies with larger data sets would reveal the changes occurring exactly during this transformation at the gene level. Furthermore, based on the differential gene expression analysis using the Deseq-2 pipeline on the T-bio info server between the Glioma neural stem cells(GSCs) and Normal neural stem cells(NSCs) samples we scrutinized 1436 significantly differentially expressed genes between GSCs and NSCs[(padj. value <0.05, log2 fold change (>=+/-1.5)]. Among them, 619 genes were found to be significantly upregulated (p.adj value <0.05, Fold change >= +1.5) genes and 817 genes were found to be significantly downregulated (p.adj value <0.05, Fold change <= −1.5) in GSCs in comparison to NSCs. Later, the dendrograms, heat map, and Hierarchical clustering revealed the significance of significantly expressed genes between GSCs and NSCs samples. A heat map generated from these genes showed that some of the genes were downregulated in GSCs samples whereas some of them were up-regulated in NSCs samples, highlighting differential expression in both populations.

Gene ontology analysis performed using Enrichr based on significant gene sets showed obvious involvement of genes in pathways like Axon growth curve pathway, molecular nervous system development pathway, amyloid beta-binding pathways, some of the amino acids metabolism pathway, Axon guidance, Extracellular Matrix organization, Collagen binding pathways, and receptor-ligand binding pathways. Furthermore, one of the most affected pathways as per our KEGG pathway analysis was the serine,threonine and glycine metabolism pathway. The associated genes with this pathway are *MAOA, MAOB, GATM, GLDC, AMT, and SHMT1*[as per Enrichr] as the key features contributing to the disturbed processes of Glycine, threonine, and serine amino acid metabolism. One of the main functions of the MAOA gene is to provide instructions for making an enzyme called monoamine oxidase A(30,31). Monoamine oxidase A breaks down molecules called monoamine by a process known as oxidation(32). *MAOA* usually is involved in the breakdown of the neurotransmitters serotonin, epinephrine, norepinephrine, and dopamine(33). Studies have shown that low-levels of dopamine in tumor tissues leads to excess stress condition, however the underlying reason still remains unknown(34). However our study depicts down-regulation of dopamine breaking gene MAOA which can be a possible reason for lower levels of dopamine in GBM. Incomplete breakdown of dopamine in this study, confirms the symptoms of GBM like hallucinations, depression, anxiety and delusions. One of the main function of *MAOB* gene is to the oxidative deamination of biogenic and xenobiotic amines. Apart from this, in association with nervous system it plays an important role in in the metabolism of neuroactive and vasoactive amines in the central nervous system and peripheral tissues(35)Neuroactive amines can .act as neurotransmitters, neuromodulators, or neurohormones(36).Biogenic amines further important biological processes like cardiovascular control, circadian rhythms, emotions, as well as learning and memory(36). Our study here elucidates that down-regulation of *MAOB* gene could be a possible reason for disturbed processes of endocrine secretion, learning and memory in GBM. *GATM* gene plays an important role in catalysing the biosynthesis of guanidinoacetate and also in the nervous system development(37).In any biologically system, poor organizational development may give rise to unwanted cells and ultimately leads to the formation of tumors, which is the possible reason for GBM formation here. Down-regulation of *GATM* gene is a vital factor playing a role in giving rise to GBM tumor formation since it depicts disturbed nervous system development.The main function of *GLDC* gene is providing instructions for making an enzyme called glycine dehydrogenase(38). Down-regulation of the GLDC gene itself portrays here the reason for disturbed metabolism of glycine amino acid in GBM. *SHMT1* is a protein coding gene responsible for glucose energy metabolism pathways.Studies have shown known association of *SHMT1* gene with leukemia(39).However down-regulation of *SHMT1* could possibly contribute to the aggressiveness of GBM. Since one of the main functions of biogenic amino acids is to support cells to make other cells, cell growth and also in DNA replication. Our study clearly depicts the dysregu;ation of crucial amino acids like Glycine,serine and threonine, which portrays a definite cause of GBM.

Apart from this, our study also portrays one of the most affected cellular processes that is the axonal growth cone. Axonal growth cone is responsible for determination of cell growth direction, extracellular motility(ECM), and provides growth direction of the axon in that direction(40). Studies have shown that disturbed ECM processes lead to invasion and motility in GBM(41).Since glial cells and axons in GBM lead to aggressive growth which ultimately to lethal tumor formation, could state lack of growth directions from axonal growth cone processes due to the down-regulation of this particular process in GBM. However investigation of the axonal growth cone pathway in association of GBM could help us gain more insights in progression of this horrendous disease.In association with this, drawing a conclusion to association of beta-amyloid protein binding pathways and it’s down-regulation in GBM, here reveal new insights in progression of GBM.

Conclusively, our study revealed significant differences in gene expression between Glioma Neural stem cells and Normal neural stem cells samples. Importantly, we confirm and validate the significant down-regulation of 6 amino acid(serine,glycine and threonine) metabolism pathway-associated genes in GSCs in comparison to NSCs samples. This most likely reveals an important clue to the etiology of this fatal brain tumor.

## Future Directions

In future studies, may be accessing the precise role of protein coding genes in relation with Glioblastoma multiforme could help us achieve therapeutic targets for GBM.However limited number of samples, was a limitation of this study, which could be overcomed with larger cohort datasets of GBM samples. Also, analysing the role of genes like *COBL, FLRT3 and GPM6A* which contribute to the significantly downregulated disturbing process of the axonal growth curve in humans, could help us achieve greater heights in treating this fatal brain tumor.

## Supplementary Data

Supplementary File 1: Supplementary Figures.

Supplementary File 2: Supplementary Tables.

## Abbreviations

GBM: Glioblastoma Multiforme
GSCs: Glioma Neural stem cells
NSCs: Normal neural stem cells
CSCs: Cancer stem cells
PCA: Principal component analysis
ECM: Extra-cellular Motility

