## Supplementary File 1 for "Gene Expression Analysis of Glioma Neural Stem Cells Shows Disturbed Amino Acids Metabolism and Axonal Growth Cone Dynamics in Glioblastoma Multiforme"

Rutvi Vaja*^1^

^1^ Navrachana  University,Vadodara,Gujarat,India

*

**Emails of Authors:**

**Correspondence:**

*Rutvi Vaja

School of Science, Department of Biomedical Sciences, Navrachana University, Vadodara, Gujarat, India

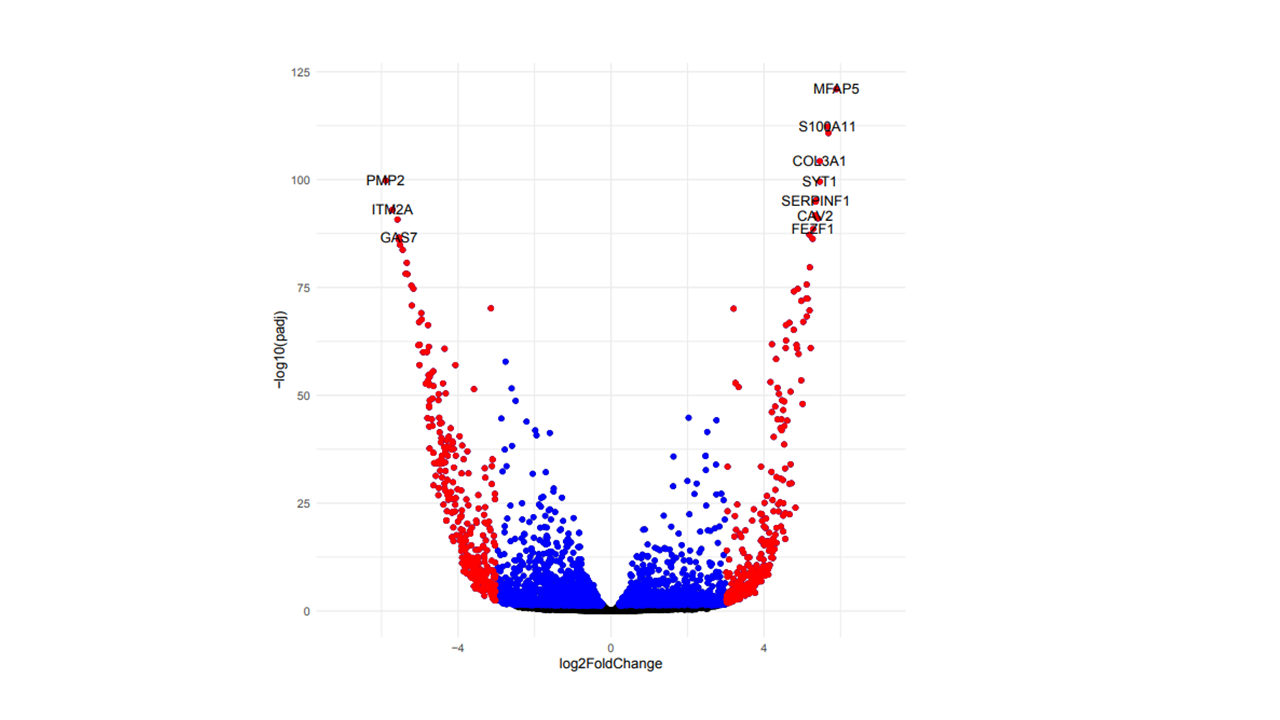

**Supplementary Figure S1: Volcano-Plot of diffrentially expressed Genes**
